# Sorting Nexin 27 (SNX27): A Novel Regulator of Cystic Fibrosis Transmembrane Conductance Regulator (CFTR) Trafficking

**DOI:** 10.1101/304717

**Authors:** Mark I. McDermott, William R. Thelin, Yun Chen, Patrick T. Lyons, Gabrielle Reilly, Martina Gentzsch, Cai Lei, Wanjin Hong, M. Jackson Stutts, Martin P. Playford, Vytas A. Bankaitis

## Abstract

The underlying defect in cystic fibrosis is mutation of the cystic fibrosis transmembrane conductance regulator (CFTR), a cAMP-activated chloride channel expressed at the apical surface of lung epithelia. In addition to its export and maintenance at the cell surface, CFTR regulation involves repeated cycles of transport through the endosomal trafficking system, including endocytosis and recycling. Many of the known disease mutations cause CFTR intracellular trafficking defects that result in failure of ion channel delivery to the apical plasma membrane. Corrective maneuvers directed at improving transport to the plasma membrane are thwarted by rapid internalization and degradation of the mutant CFTR proteins. The molecular mechanisms involved in these processes are not completely understood but may involve protein-protein interactions with the C-terminal type I PDZ-binding motif of CFTR. Using a proteomic approach, we identify sorting nexin 27 (SNX27) as a novel CFTR binding partner in human airway epithelial Calu-3 cells. SNX27 and CFTR interact directly, with the SNX27 PDZ domain being both necessary and sufficient for this interaction. SNX27 co-localizes with internalized CFTR at sub-apical endosomal sites in polarized Calu-3 cells, and either knockdown of the endogenous SNX27, or over-expression of a dominant-negative SNX27 mutant, resulted in significant decreases in cell surface CFTR levels. CFTR internalization was not affected by SNX27 knockdown, but defects were observed in the recycling arm of CFTR trafficking through the endosomal system. Furthermore, knockdown of SNX27 in Calu-3 cells resulted in significant decreases in CFTR protein levels, consistent with degradation of the internalized pool. These data identify SNX27 as a physiologically significant regulator of CFTR trafficking and homeostasis in epithelial cells.

## Introduction

Cystic fibrosis (CF) is caused by mutations in the gene encoding the cystic fibrosis transmembrane conductance regulator (CFTR), a cyclic AMP-dependent chloride ion channel localized to the apical membrane of secretory epithelia (Riordan, 2008; Riordan et al., 1989). Of the more than 1400 identified CFTR mutations, the most prevalent mutation involves deletion of phenylalanine at position 508 (ΔF508). This mutation causes protein misfolding, retention of the malfolded protein in the ER, and subsequent degradation by the proteosome via the ER-associated degradation pathway (ERAD) (Chang et al., 1999).

The ΔF508 mutation allele is present in a homozygous state in approximately 70% of CF patients (Tsui, 1992). Given the prevalence of this disease allele, major therapeutic efforts have been trained on strategies for enabling CFTR to escape the ER and be delivered to the apical plasma membrane (Becq, 2010). Chemical or temperature-shift maneuvers that “chase” CFTR out of the ER increase functional CFTR at the plasma membrane in cell culture systems (Brown et al., 1996; Denning et al., 1992; Egan et al., 2004; Grubb et al., 2006). However, the benefits realized by such strategies are largely negated by rapid clearance of the “rescued” ΔF508 CFTR from the cell surface with subsequent degradation (Swiatecka-Urban et al., 2005). The rate of wild type cell surface CFTR pool internalization is approximately 3-4% per minute (Barriere et al., 2011; Swiatecka-Urban et al., 2004). Given that the half-life of CFTR is approximately 16 h, the internalized CFTR protein must normally be recycled back to the cell surface efficiently rather than be sorted to the lysosome for degradation (Lukacs et al., 1994; Sharma et al., 2004; Ward and Kopito, 1994). Although CFTR traffics through early and recycling endosomal compartments, the mechanisms controlling CFTR recycling are not completely understood (Gentzsch et al., 2004; Picciano et al., 2003). For example, Rme1 and syntaxin 16 are both proposed to regulate CFTR recycling, but how these do so remains unclear (Gee et al., 2010; Picciano et al., 2003).

Protein-protein interactions regulate nearly all aspects of CFTR biology (Balch and Yates, 2011; Gee et al., 2011; Koulov et al., 2010; Swiatecka-Urban et al., 2004; Wang et al., 2006). The extreme C-terminus of CFTR contains a class I PDZ binding motif (DTRL) proposed to play important roles in the unconventional plasma membrane secretion, membrane mobility, cell surface stability, and cAMP-dependent channel activation of CFTR (Bates et al., 2006; Gee et al., 2011; Haggie et al., 2004; Moyer et al., 1999; Short et al., 1998; Swiatecka-Urban et al., 2002) Several PDZ-domain containing proteins bind the CFTR C-terminus. These include members of the sodium hydrogen exchange regulatory factor (NHERF) family, the Golgi-associated PDZ and coiled-coil motif containing protein GOPC (also known as the CFTR-associated ligand or CAL), and SHANK2 (Cheng et al., 2002; Han et al., 2006; Moyer et al., 1999; Short et al., 1998; Sun et al., 2000; Wang et al., 1998). NHERFs contain multiple PDZ domains and localize to the apical membrane of epithelial cells (Short et al., 1998; Sun et al., 2000). The observation that NHERFs can interact simultaneously with CFTR and the actin cytoskeleton suggests that NHERFs control CFTR retention and activity at the cell surface (Weinman et al., 2006). Consistent with this idea, NHERF-1 (solute carrier family 9 (sodium/hydrogen exchanger) isoform 3 regulatory factor 1 or SLC9A3R1) and NHERF-2 (solute carrier family 9 (sodium/hydrogen exchanger) isoform 3 regulatory factor 2 or SLC9A3R2) both stimulate CFTR channel activity without any effect on the CFTR intracellular trafficking itinerary (Benharouga et al., 2003; Raghuram et al., 2001; Wang et al., 2000).

In contrast to the NHERF proteins, GOPC contains a single PDZ domain and is located primarily on the Golgi apparatus (Cheng et al., 2002). Hence, GOPC is proposed to regulate initial CFTR trafficking from a post-ER compartment to the apical plasma membrane. GOPC negatively regulates CFTR expression at the cell surface and may compete with NHERF for binding to the CFTR C-terminus (Cheng et al., 2002). A second negative regulator of CFTR channel activity, SHANK2, is localized to the apical region of epithelial cells (Kim et al., 2004; Lee et al., 2007). Kinetic analyses reveal that the dissociation constants (K_d_) of SHANK2 and NHERF-1 from the PDZ-binding motif of CFTR are similar and that, like GOPC, both SHANK2 and NHERF-1 compete for the same binding site on CFTR (Lee et al., 2007). What role SHANK2 plays in CFTR trafficking, if any, is unknown.

Herein, we identify the PDZ-containing protein sorting nexin (SNX27) as a novel CFTR binding partner. SNX27 localizes to both early and recycling endosomes, and promotes CFTR recycling from these compartments to the cell surface. These data identify SNX27 as a key regulator of CFTR trafficking and protein levels in epithelial cells -- a function consistent with previous demonstrations that SNX27 controls endosomal trafficking of other ion channels and G-protein coupled receptors (Lauffer et al., 2010; Lunn et al., 2007; Temkin et al., 2011).

## Materials and Methods

### Animal use and ethics statement

The experimental procedures used in this study strictly followed the U.S. Public Health Service Policy of Humane Care and Use of Laboratory Animals and were approved by the Institutional Animal Care and Use Committee at the University of North Carolina at Chapel Hill (IACUC ID:05-050.0).

### Affinity purification and mass spectrometry

Mouse kidneys were obtained from wild-type animals using approved protocols and homogenized in binding buffer 150 (BB150: 50 mM Tris pH 7.6, 150 mM NaCl, 0.2% CHAPS, 10 mM EDTA) plus Complete™ protease inhibitor cocktail (Roche Applied Science, Illinois, USA) and 1 mM PMSF on ice. Lysates were incubated at 4°C for 1 h prior to ultracentrifugation at 100,000 × g for 1 h. Supernatants were pre-cleared with streptavidin agarose beads (Sigma-Aldrich, St Louis, Missouri, USA) and incubated with wild type CFTR or mutant affinity matrices for 4 h at 4°C (20 nmoles of wild type or mutant CFTR peptide bound to 100 μl of streptavidin beads). The wild type peptide corresponded to the last 10 amino acids of CFTR (1471-1480), and mutant peptide (1471-1480/4G) substituted the D-T-R-L PDZ-binding motif residues to glycines. Fractions bound to the matrices were washed five times in BB150 buffer, washed once in 10 mM sodium phosphate pH 7.5 plus 75 mM NaCl and eluted in 10% formic acid. The formic acid concentration was then adjusted to 1% with ddH2O, and the samples were lyophilized prior to processing and analysis by LC-MS/MS as previously described (Thelin et al., 2005) using a Waters Q-tof micro, hybrid quadropole orthogonal acceleration time-flight mass spectrometer (Waters, Manchester, UK). Western blot analysis, to confirm interactions, was performed using the Odyssey infrared imaging system (LI-COR, Lincoln, Nebraska, USA).

### Antibodies and other reagents

CFTR monoclonal antibodies 217, 293, 570, and 596 were kindly provided by Dr John R. Riordan (University of North Carolina, Chapel Hill, USA). Antiserum generated against NHERF-1 was obtained from Affinity bioreagents (Rockford, Illinois, USA); antiserum against NHERF-3 kindly provided by David Silver (Columbia University, New York, USA) and antiserum against ezrin was obtained from Santa Cruz Biotechnology (Santa Cruz, California, USA). The polyclonal SNX27 antibody used in this study has been previously described (Rincon et al., 2007). Detection of HA-tagged extope-CFTR by immunofluorescence and western blotting was performed using HA.11 (Covance, Princeton, New Jersey, USA). Secondary anti mouse HRP (abcam) was used to detect antibodies against endogenous CFTR or the HA-tag of extope CFTR 1:10,000 dilution, blots were exposed to film and densitometry performed using an Imagequant™ LAS 4000 (GE). Detection of Myc-tagged proteins by immunofluorescence, immunoprecipitation, and western blotting was performed using anti-Myc 9E10 antibody (Covance). Secondary anti-rabbit antibody conjugated with IRDye 800CW or antimouse conjugated with IRDye 680 were purchased from LI-COR and used for western blotting detection at 1:10000 dilution. Secondary antibodies used for immunofluorescent detection were anti-rabbit Alexa Fluor 488 and anti-mouse Alexa Fluor 594 from Invitrogen/Molecular Probes (Carlsbad, California, USA) and used at a 1:1000 dilution. All chemicals unless otherwise indicated were obtained from Sigma.

### Cell culture and transfections

Baby Hamster Kidney cells (BHK-21), HeLa, and Calu-3 cells were obtained from the American Type Culture Collection (ATCC, Manassas, Virginia, USA). BHK-21 and HeLa cells were cultured in high glucose, Dulbecco’s Modified Eagle Medium (Invitrogen) supplemented with 10% fetal bovine serum (Sigma) and maintained in humidified chambers with 5% CO_2_ at 37°C. Calu-3 cells were cultured as previously described (Thelin et al., 2005). Cells were transiently transfected using Fugene 6 (Roche), Lipofectamine LTX (Invitrogen), or Lipofectamine 2000 (Invitrogen) as per the manufacturer’s instructions. Stable BHK-21 cell lines expressing extope-CFTR WT were maintained as previously described (Gentzsch et al., 2004).

### Determination of mammalian cell number, size and viability

Post trypsinization cells were resuspended in cell culture media and mixed with an equal volume of trypan blue (Invitrogen), prior to being analyzed using a Countess^TM^ automated cell counter (Invitrogen) according to the manufacturers instructions. Viability is provided as the % of live cells within the count. Cell size is provided as the average diameter of the trypsinized cells (μm). At least 100 cells were counted per data point, all experiments were performed in quadruplet.

### Cloning and mutagenesis of SNX27

The cDNA sequence of human SNX27b was amplified by PCR and cloned into the *Bam*HI and *Xho*I sites of pCMV-Myc (Invitrogen), PET28c (Novagen, EMD chemicals, Gibbstown, New Jersey, USA), and pEGFP (Clontech, Mountain View, California, USA). SNX27b R196G and Y53A/F55G mutations, within the PX and PDZ domains, respectively, were generated using the QuikChange^®^ Site-Directed Mutagenesis kit (Stratagene, Santa Clara, California, USA). SNX27b mutants lacking the PDZ and FERM-like (amino acids 274-360) domains were generated by inverse PCR. DNA sequencing was performed to confirm all constructs were generated without undesired secondary mutations.

### *In vitro* binding assays

Binding assays were performed essentially as described previously (Thelin *et al*., 2005). Briefly 5 μg of the indicated peptides were immobilized to streptavidin beads (Sigma) in BB150 buffer at 4 °C, and washed three times with BB150 to remove unbound peptide. BHK cells, transfected with wild type or mutant SNX27, were lysed in BB150 containing protease inhibitors (Roche) by passage through a 27-gauge syringe at 4 °C. Lysates were incubated with the streptavidin conjugated peptides or uncoated control beads for 2 h, washed three times with BB150, and bound proteins eluted by boiling in Laemmli buffer (62.5 mM Tris-HCl, 2% SDS, 25% glycerol, and 0.01% Bromophenol Blue pH 6.8). Samples were analyzed by SDS-PAGE and western blotting using the Odyssey infrared imaging system (LI-COR™).

### Generation of glutathione-S-transferase (GST)-SNX27 fusion proteins

The PDZ domain of SNX27 (43-136) was amplified from SNX27b using PCR, cloned into the *Bam*HI and *Eco*RI sites of pGEX2T (GE Lifesciences, Piscataway, New Jersey, USA) and transformed into BL21 (DE3) cells (Stratagene). SNX27-PDZ fusion protein expression was induced with 0.1 mM IPTG (isopropyl-b-D-thiogalactopyroside) for 3 h at room temperature. The bacterial cells were lysed by sonication for 1 min in PBS and proteins solubilized in Triton X-100 diluted to 1% final volume. Fusion proteins were collected by incubation at 4°C with glutathione-Sepharose 4B beads (GE Life Sciences).

### GST-pulldowns

To establish if the SNX27 PDZ domain is sufficient for CFTR interaction, lysates from BHK, BHK-extope CFTR and CALU-3 cells prepared as above were incubated with either GST alone or a fusion of the PDZ domain of SNX27 with GST (GST-PDZ). Bound proteins were resolved by SDS PAGE and detected using western blotting for CFTR as described above.

### Immunofluorescence of CFTR and SNX27 in BHK and HeLa cells

BHK cells were plated on fibronectin (10 μg/ml)-coated glass coverslips and transfected as indicated. Cells were fixed with 4% paraformaldehyde (PFA) for 15 min, permeabilized with 0.1% Triton X-100 for 15 min, and incubated in blocking buffer (2% bovine serum albumin (BSA) in PBS) for 1 h at 37°C. The fixed coverslips were incubated with HA.11 antibody at 1:500 and anti-myc 9E10 at a 1:500 dilution (to detect CFTR and SNX27, respectively) in blocking buffer for 1h at 37°C. Coverslips were washed extensively with PBS and incubated with secondary antibody in blocking buffer for 1 h. Following further washing with PBS, coverslips were mounted using FluorSave™ (Calbiochem). Confocal images were acquired at room temperature using a Zeiss LSM 510 system mounted on a Zeiss Axiovert 100M microscope with an oil immersion Plan-Apochromat 63x/1.4 DIC objective lens (Carl Zeiss Microscopy, Thornwood, New York, USA). Excitation wavelengths of 488 nm (10%), 543 nm (100%), and 364 nm (5%) were used for detection of Alexa Fluor 488, Alexa Fluor 594 and DAPI, respectively. Fluorescent emissions were collected in filters BP505-550 nm, LP560 nm and BP385-470 nm, respectively. All pinholes were set at 1.00 Airy units (AU), which corresponds to an optical slice of 0.8 *μ*m. All confocal images were of frame size 512 pixels by 512 pixels, scan zoom of 2 and line averaged 4 times. All images were analyzed using the Zeiss AIM software version 3.2 sp2 (Carl Zeiss GmbH, Heidelberg, Germany).

### Immunofluorescence imaging in polarized Calu-3 cells

Cells were grown on permeable supports and studied after achieving transepithelial resistance (Rt)> 300 Ωcm^2^. Cells were fixed with ice-cold methanol for 2 min and blocked for 2 h in PBS containing 2 mg/ml BSA, 4% non-fat milk, 1% fish gelatin and 0.1% Triton X-100. Primary antibodies were diluted in the same buffer and incubated at 4°C overnight. Cells were washed extensively in buffer lacking milk and incubated with secondary antibodies as described above for 1 h, prior to washing and mounting.

### Imaging of CFTR internalization

To track the internalization of CFTR, BHK cells over-expressing wild type extope-CFTR and either GFP-SNX27 or GFP-SNX27 ΔPDZ were cultured on fibronectin-coated glass cover slips. Live cells were chilled to 4°C on ice and incubated with HA-antibody as previously described (Gentzsch et al., 2004). Unbound antibody was removed by washing with PBS and the cells returned to 37°C in growth media for various times as indicated, to allow the CFTR-antibody-complexes to internalize. Cells were fixed in 4% PFA, incubated in blocking solution, permeabilized with 0.1% Triton X-100 in PBS and visualized with secondary antibodies as described above.

### Generation of SNX27 knockdown cell lines

Stable HeLa and Calu3 cell lines expressing SNX27 silencing or non-silencing (control) shRNAs were generated using the pINDUCER10 inducible lentiviral shRNA system (Meerbrey *et al*., 2011). Briefly, SNX27 was subcloned into the *MluI* and *XhoI* sites of pINDUCER10. Lentiviral supernatants were obtained by transiently transfecting HEK293T cells using Lipofectamine LTX (Life Technologies). Control lentivirus was generated using an otherwise identical method but by substituting a control firefly luciferase plasmid. Stable knockdown HeLA and Calu3 cell lines were then generated by viral transduction and selected and maintained with puromycin as previously described (Meerbrey et al., 2011). shRNA expression was induced in the stable cell lines by addition of 1 μg/ml doxycycline for 72 h prior to each experiment.

### Quantification of cell surface CFTR by laser scanning cytometry

Surface CFTR levels were measured in BHK-extope CFTR stable cells transfected with the indicated GFP-tagged SNX27 constructs. Cells were cultured on fibronectin-coated glass cover slips. Live cells were chilled to 4°C on ice and incubated with HA-antibody as previously described (Gentzsch et al., 2004). Unbound antibody was removed by washing with PBS. Cells were then fixed in 4% PFA, incubated in blocking buffer containing a secondary anti-mouse Cy5 antibody (Jackson Immunoresearch, West Grove, Pennsylvania, USA), washed and mounted as described above. Cell surface CFTR was measured using an iCys^®^ Research Imaging Cytometer (CompuCyte; Cambridge, MA, USA). Slides were observed with an attached IX71 Olympus microscope and DONPISHA XC-003 3CCD Color Vision Camera module (Olympus, Center Valley, Pennsylvania, USA). The following standard filter setting was used for detection: Green filter (530/30 FITC) and Long Red filter (650/LP Cy5). Excitation was performed with a 488 nm argon laser and a 633 nm helium-neon laser. Two scan areas were selected for each slide and were scanned using a 40x objective.

### Single particle trafficking of CFTR

Single particle tracking (SPT) experiments were performed similar to those previously described (Chen et al., 2009). Briefly, BHK-extope-CFTR cells were transiently transfected with GFP, GFP-SNX27, or GFP-SNX27 lacking the PDZ-binding domain (ΔPDZ). Cell surface CFTR was labeled in these cells, on ice, using a biotinylated antiHA antibody (Covance), followed by anti-biotin-conjugated 20 nm-diameter quantum dots 605 (Invitrogen). Cells were returned to warmed media (37°C) and quantum dot-labeled CFTR was imaged using an Olympus IX71 microscope in the red fluorescence channel for 100 frames/particle at 1 Hz. ((van der Schaar et al., 2008); (Kulkarni et al., 2006) (Murray et al., 2000; van Weering et al., 2012). Experiments were performed in triplicate. For each experimental condition at least 30 particles from a total of 5 cells, were recorded over the course of 3 experiments. Movies were analyzed using Metamorph software (Universal Imaging, Burbank, California, USA), and all trajectories were visually inspected to ensure correct tracking.

## Results

### Identification of SNX27 as a candidate CFTR binding partner

The C-terminal PDZ-binding motif of CFTR interacts with several PDZ-domain containing proteins with emerging roles in CFTR regulation (Cheng et al., 2002; Han et al., 2006; Moyer et al., 1999; Short et al., 1998; Sun et al., 2000; Wang et al., 1998). In order to identify novel proteins interacting with this region, synthetic biotinylated peptides corresponding to the last 10 amino acids of CFTR (1471-1480) containing the C-terminal PDZ-binding motif (DTRL), or control peptides (1471-1480/4G) where DTRL is substituted with four glycine residues, were synthesized and immobilized on streptavidin agarose beads (Fig. 1A). The bead-peptide complexes were used as affinity ligands with lysates from mouse kidney tissue. Bound proteins were eluted with formic acid and subsequently digested in-solution with trypsin. Peptide analysis by nano-liquid chromatography tandem mass spectrometry (nano-LC MS/MS) identified candidate CFTR binding partners.

**Fig. 1.**
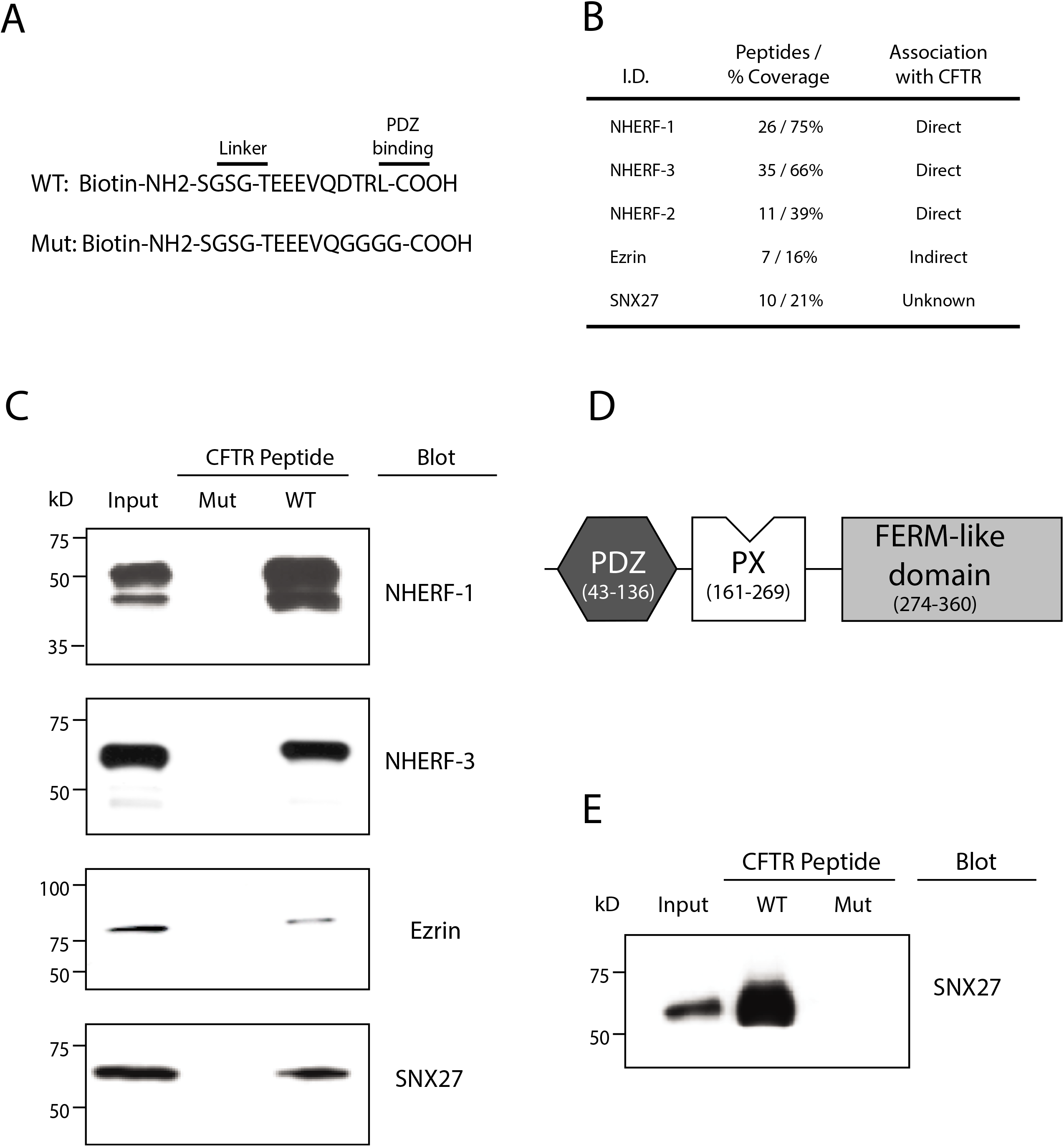
Identification of interacting proteins with the CFTR C-terminus. Sequences of synthetic biotinylated C-terminal CFTR peptides used in this study. Wild type CFTR C-terminal peptide contained the type I PDZ binding motif (DTRL), which was replaced with GGGG for mutant control (A). Summary of proteins bound to wild type, but not mutant C-terminal CFTR peptide (B). Biotinylated wild type and mutant peptides were immobilized on streptavidin beads and incubated with lysates from mouse kidney tissue. Bound proteins were analyzed by western blotting using antisera against NHERF-1, NHERF-3, ezrin and SNX27 respectively. All proteins associate specifically with the wild type but not the mutant CFTR C-terminal peptide (C). SNX27 was identified as a novel CFTR interacting protein. SNX27 is a member of the PX domain containing sorting nexin family and is unique in possessing a PDZ and a FERM-like domain (D). Purified His-tagged SNX27 was incubated with the indicated CFTR peptides. Peptide-bound SNX27 was identified by western blotting (E).

Propionyl coenzyme A carboxylase (PCC) was identified as an expected contaminant in the streptavidin-bound fraction. This enzyme is an endogenous biotin modified protein (Wood and Barden, 1977), and its association with streptavidin agarose beads was peptide independent (supplemental figure). Other expected binding partners were also recovered. These included members of the NHERF family previously shown to bind the CFTR C-terminus, and ezrin which complexes with CFTR indirectly via its ability to bind NHERF-1 and −2. One novel CFTR interactor was detected in these experiments, however, and it was identified as the PDZ-domain-containing protein sorting nexin 27 (SNX27) (Fig. 1B and supplemental figure 1). SNX27, like the NHERF proteins and ezrin, satisfied the specific binding criterion as all these proteins failed to bind control peptides (Fig. 1 and see below).

### SNX27 interacts with the CFTR C-terminus

In support of the MS data, peptide affinity purifications followed by immunoblotting demonstrated that SNX27 is extracted from murine kidney cell lysates by WT (1471-1480) but not mutant (1471-1480/4G) CFTR bait peptides. As expected, the previously known CFTR interactors: NHERF-1, NHERF-3, and ezrin also bound to specifically to WT CFTR peptide (Fig. 1C). SNX27 belongs to a 33-member sorting nexin protein family implicated in protein trafficking. Like the other family members, SNX27 has a PX (phosphoinositide-binding phox homology) domain but is unique in possessing a PDZ domain and a putative FERM-like domain (FLD) (Fig. 1D). To establish whether SNX27 bound CFTR directly, purified His-tagged SNX27 was incubated with CFTR C-terminal peptides. Immunoblotting of precipitates demonstrated clear and specific binding of SNX27 to WT CFTR peptide (Fig. 1E).

To determine whether SNX27 associates with CFTR in human airway epithelial cells, endogenous CFTR was precipitated from Calu-3 cell lysates, and presence of SNX27 in those immune complexes was interrogated by immunoblotting (Fig. 2A). SNX27 co-precipitated with CFTR, but was not detected in control IgG immune complexes (Fig. 2A, middle panel). The specificity of CFTR/SNX27 interaction was further indicated by the absence of other related sorting nexins in CFTR immunocomplexes (e.g. SNX2; Fig. 2A bottom panel).

**Fig. 2.**
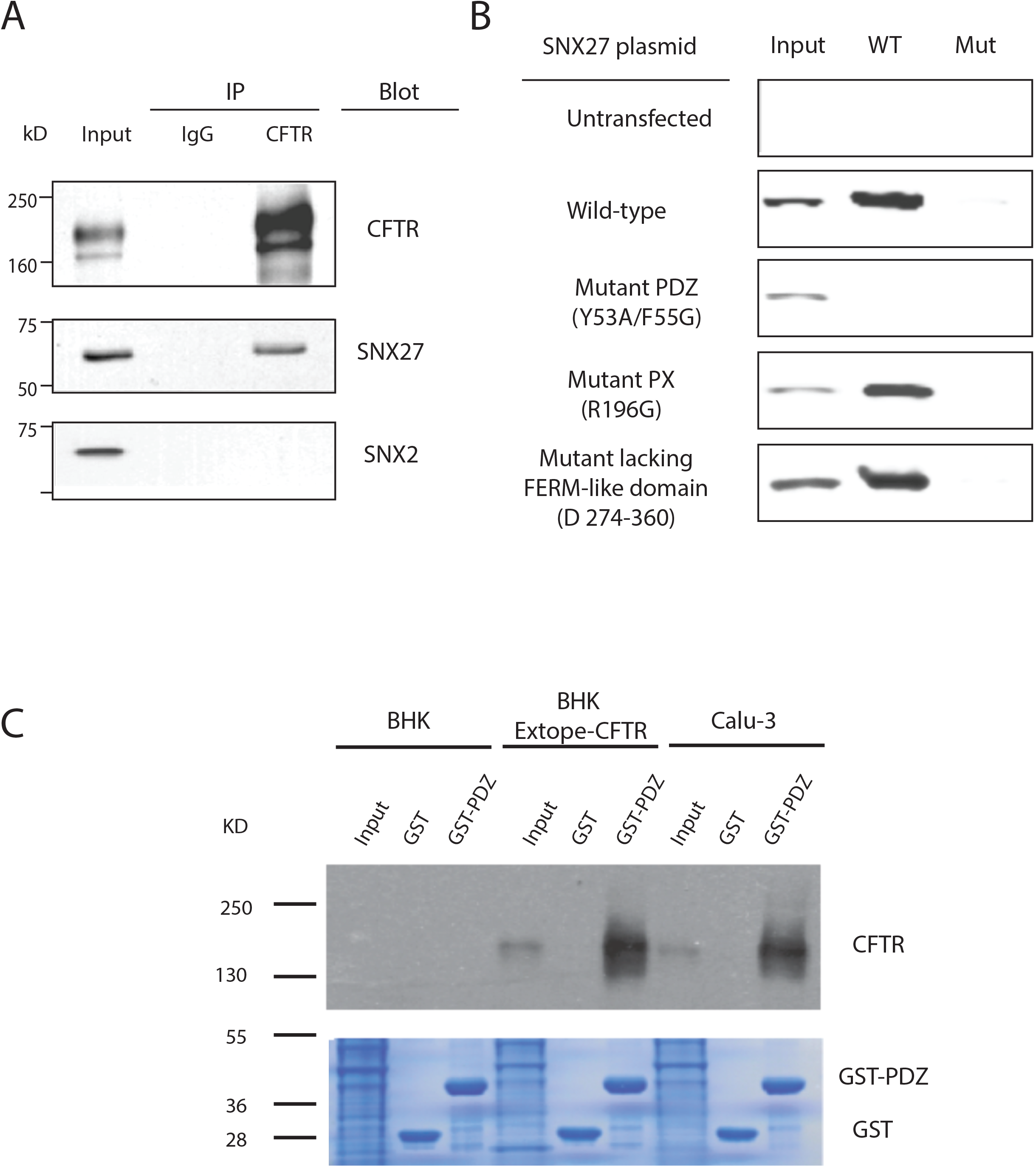
Interaction between SNX27 with CFTR occurs endogenously in lung epithelial cells and is PDZ dependent. Endogenous CFTR was immunoprecipitated from Calu-3 cell lysates and analyzed by western blotting using antibodies against SNX27, SNX2 or CFTR. SNX27 but not SNX2 co-immunoprecipitated with CFTR (A). Inputs 5% of lysate used for the IP. Cell lysates were prepared from BHK cells transfected with the indicated myc-tagged SNX27 constructs. Lysates were incubated with the biotinylated wild type or mutant CFTR peptides immobilized on streptavidin beads. Interacting myc-tagged SNX27 proteins were identified by western blotting for myc (B). Lysates from BHK, BHK-extope-CFTR or Calu-3 cells were incubated with GST-SNX27 or control GST proteins. GST-bound proteins were analyzed by western blotting using an antibody specific for CFTR (C).

The SNX27 PDZ-domain was an obvious candidate motif for controlling interaction of this sorting nexin with CFTR. To test this possibility, a Myc-tagged SNX27 PDZ domain variant was generated where the conserved GYGF motif of the hydrophobic PDZ ligand-binding pocket was converted to GGGG (mPDZ Y53G/F55G). Such mutation of this motif within the canonical carboxylate-binding loop is predicted to substantially decrease affinity of the PDZ domain for its ligand (Doyle et al., 1998, Shultz et al., 1998, Hiller et al., 1999; Hung and Sheng 2002). As additional control, a Myc-tagged SNX27 PX domain mutant (mPX) was generated where a critical Arg residue required for mediating interaction with lipid head groups was converted to Gly (R196G) (Lee et al., 2005; Stahelin et al., 2003). A Myc-tagged SNX27 FERM-like domain deletion mutant (ΔFLD) was also generated. WT or mutant SNX27 proteins were subsequently expressed in BHK cells, and lysates from transfected BHK cells were incubated with either WT or mutant CFTR-peptides immobilized on streptavidin beads (Fig. 2B). Bound proteins were probed by immunoblotting with myc-antibodies. WT Myc-SNX27, Myc-SNX27-ΔFLD and mPX all bound strongly and specifically to CFTR peptide, whereas Myc-SNX27-mPDZ did not bind at all (Fig. 2B).

The SNX27 PDZ domain was also sufficient to mediate CFTR binding. A recombinant chimera where the SNX27 PDZ domain was fused to GST (GST-SNX27-PDZ) was purified, immobilized on glutathione agarose beads, and incubated with: (i) native BHK cell lysates (which lack endogenous CFTR), (ii) lysates of BHK cells stably expressing extope-CFTR, and (iii) Calu-3 cell lysates. GST-SNX27-PDZ captured CFTR from BHK extope-CFTR and CALU-3 lysates. By contrast, CFTR was not captured by immobilized GST alone, or when immobilized GST-SNX27-PDZ was incubated with lysate prepared from untransfected BHK cells (Fig. 2C). These collective data demonstrate a direct and specific interaction between the C-terminal CFTR PDZ-binding motif and the PDZ domain of SNX27 in human airway epithelial cells.

### SNX27 localization is consistent with a role in the endosomal recycling of CFTR

SNX27 localizes to early and recycling endosomes and belongs to a family of proteins implicated in endocytic trafficking (Rincon et al., 2007). In agreement with previous reports, we observed PX-domain dependent co-localization of SNX27 with markers for the early sorting and recycling endosomes in a number of non-polarized cell lines (Cai et al., 2011; Hayashi et al., 2012; Joubert et al., 2004; Lauffer et al., 2010; Lunn et al., 2007; Rincon et al., 2011; Rincon et al., 2007; Temkin et al., 2011). Internalized CFTR was also previously observed to co-localize with markers for the early sorting and recycling endosomes, suggesting the CFTR-SNX27 interaction is of biological relevance (Gentzsch et al., 2004; Silvis et al., 2009). To determine whether CFTR and SNX27 co-localize in cells, the cell surface pool of CFTR was labeled, and internalization and subsequent intracellular trafficking of this pool were followed. Due to the lack of an available antibody that recognizes an extracellular epitope presented by endogenous CFTR protein, we utilized the external HA-tag of exogenously expressed extope-CFTR in BHK cells. This represents a well-characterized substrate for monitoring CFTR endosomal trafficking (Gentzsch et al., 2004). Following incubation with anti-HA antibody on ice, cells were warmed, and intracellular CFTR trafficking was visualized by confocal microscopy and related to that of GFP-SNX27 (Fig. 3).

**Fig. 3.**
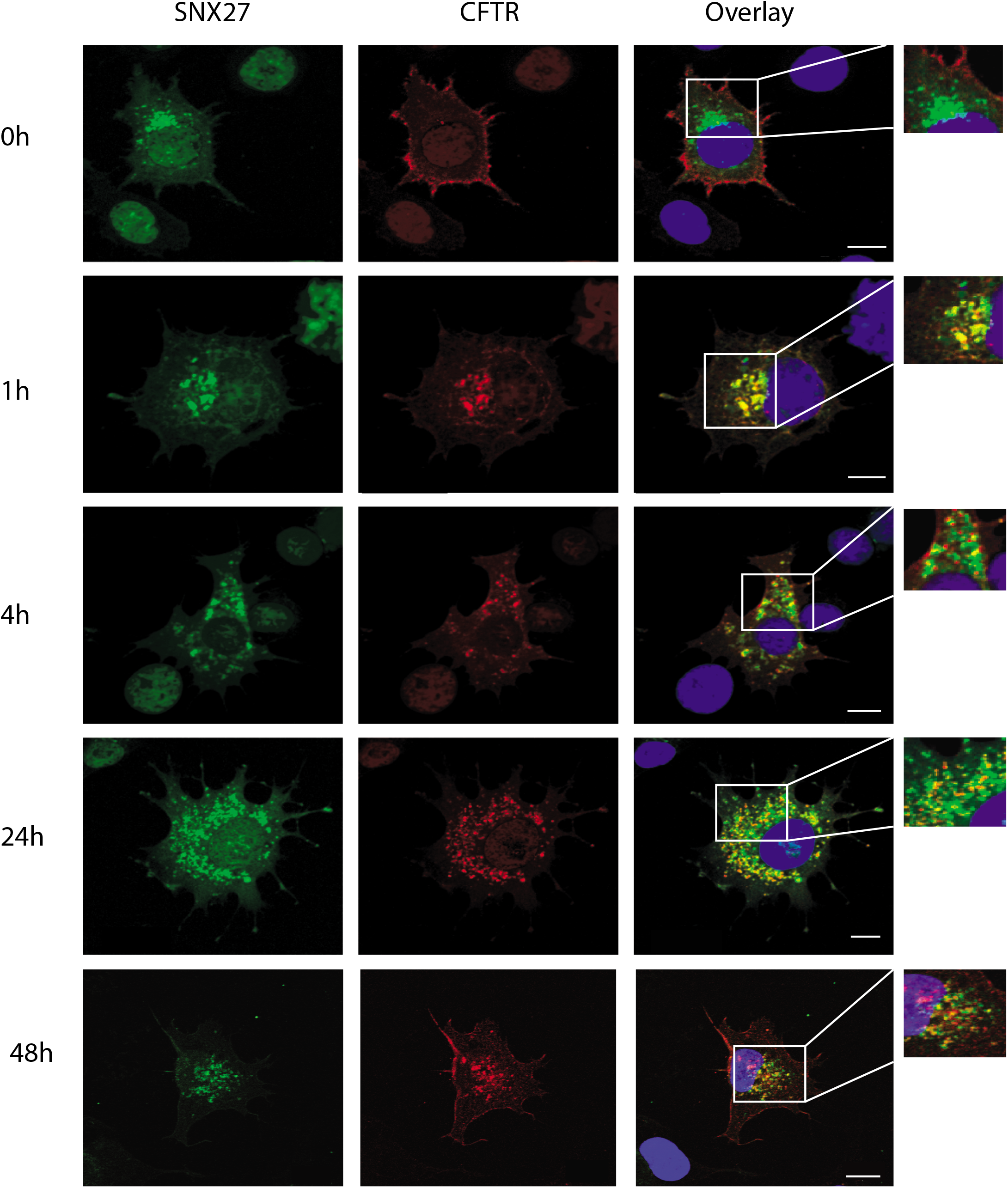
Internalized CFTR temporally co-localizes with SNX27 positive vesicles. BHK-CFTR cells were transiently transfected with GFP-SNX27. The cells were placed on ice and cell surface CFTR labeled with anti-HA antibody. The cells were subsequently restored to normal physiological conditions for the indicated times. Cells were fixed, permeabilized and incubated with secondary antibody to allow visualization of the CFTR protein. SNX27 was visualized with GFP. Scale bars, 10 μm.

Within 5 min, a CFTR pool was detected in sub-plasma membrane vesicles, with the majority of internalized protein being localized in regions clearly removed from those occupied by SNX27. By 15 and 30 min, the degree of co-localization steadily increased and, by 1 h, co-localization between the two proteins was clearly apparent in endosomal compartments (Fig. 3). While a pool of CFTR remained in SNX27 positive compartments at 4 h post-internalization, some of the internal CFTR fraction had mobilized to a distinct intracellular station lacking SNX27 (Fig. 3). By 24 h, most CFTR was localized to intracellular compartments that were smaller and more numerous than those observed at earlier time points, and were completely distinct from those harboring SNX27 (Fig. 3). By 48 h, most of the CFTR occupied distinct intracellular localizations from those of SNX27, or had recycled back to the cell surface (Fig. 3, bottom row). Enhanced SNX27 expression did not obviously alter CFTR internalization profiles (data not shown).

### SNX27 regulates CFTR levels at the plasma membrane

The trafficking data indicated the interaction of CFTR with SNX27 occurs in endosomal compartments, thereby raising the possibility that SNX27 controls CFTR recycling from endosomal compartments to the cell surface. To address whether SNX27 regulates size of the plasma membrane CFTR pool, HeLa cells were transduced with lentiviruses supporting inducible expression of either shRNA targeting SNX27 mRNA, or control shRNA. SNX27 protein levels were then assessed 72 h post induction by immunoblotting. Quantification using LI-COR^TM^ imaging reported 90% knockdown of SNX27 protein levels compared to controls (Fig. 4A). To examine how SNX27 regulates CFTR trafficking, both control and SNX27 knockdown cells were transiently transfected with extope-CFTR and cell surface CFTR was visualized by confocal microscopy using the same settings to image both sample sets. SNX27 knockdown cells displayed greatly reduced levels of cell surface CFTR compared to controls (Fig. 4B). We scored a 55% reduction in the number of cells expressing cell surface CFTR in SNX27 knockdown cells compared with control populations (Fig. 4C). Laser-scanning cytometry reported that the SNX27 knockdown population displayed a 28% reduction in cell surface CFTR when compared with control cells (Fig. 4D). Importantly, SNX27 knockdown did not affect cell viability or size (data not shown), indicating SNX27 diminution did not exert global effects on endosomal trafficking.

**Fig. 4.**
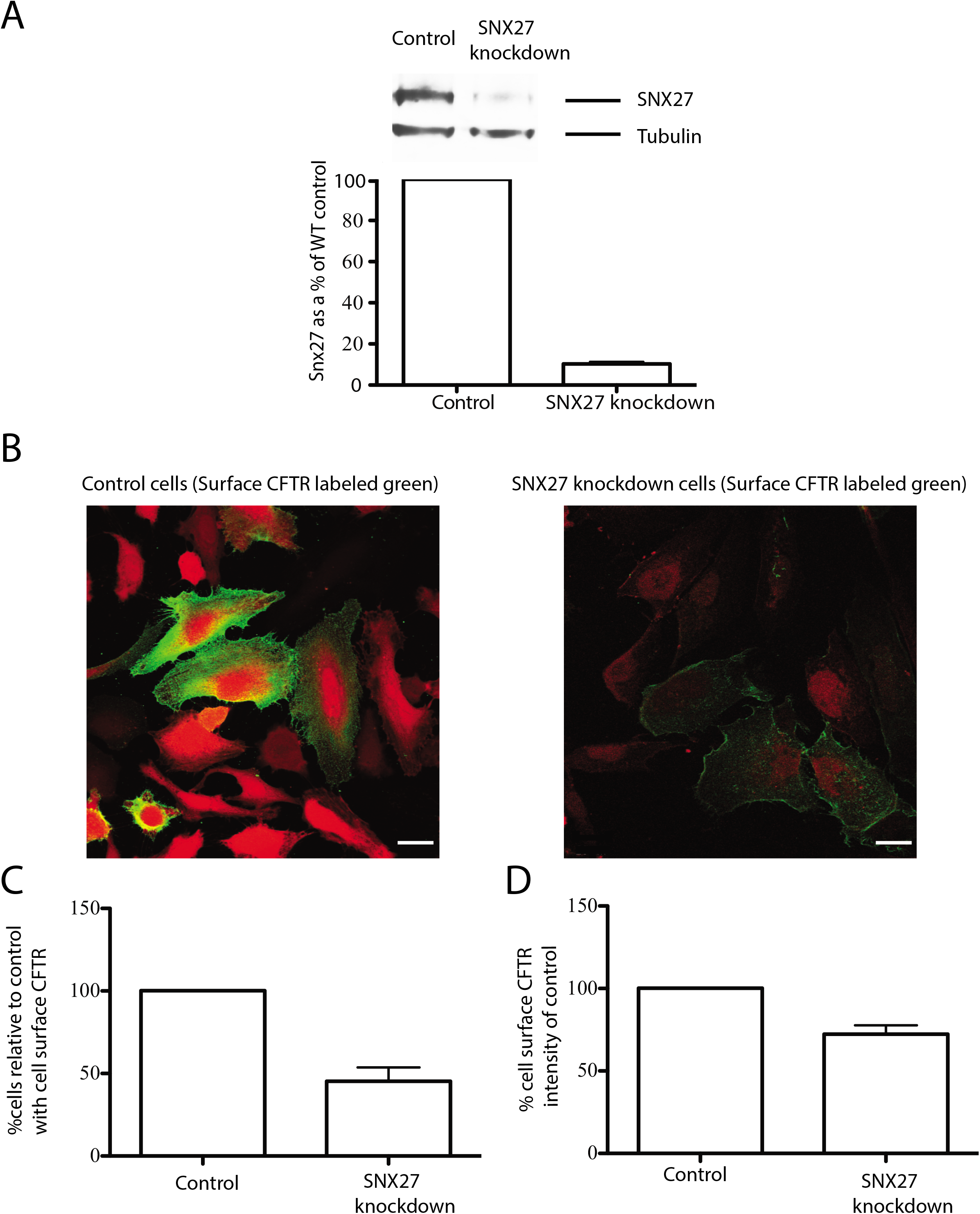
SNX27 regulates cell surface CFTR and CFTR protein levels. Stable SNX27 knockdown and control HeLa cell lines were generated using lentivirus expressing shRNA (A). Control and SNX27 knockdown cells were transiently transfected with extope-CFTR. Cells were fixed and cell surface CFTR visualized using an antibody to the extracellular HA tag (green), red fluorescence indicates presence of lentivirus. Both sets of images were taken using the same microscopy settings (note the red fluorescence is naturally lower in the knockdown versus the control cells) Scale bars, 10 μm (B). Quantification of the number of transfected cells displaying cell surface CFTR was scored by visual counting in control and SNX27 knockdown populations. At least 200 cells were counted per sample. Data was analyzed by students two-tailed t test *P* < 0.005. *n* = 4. Data shown are means +/-s.e.m. (C). Laser scanning cytometry was used to quantify overall levels of cell-surface CFTR in control versus SNX27 knockdown populations. At least 200 cells were counted per sample. Data was analyzed by students two-tailed t test. *P* < 0.005. *n* = 3 and are displayed as means +/- s.e.m. (D).

### SNX27 PDZ domain is required for regulation of CFTR trafficking

To determine the contribution of the SNX27 PDZ domain to regulation of the CFTR trafficking itinerary, BHK-extope-CFTR expressing cells were transiently transfected with GFP-SNX27 wild-type, GFP-SNX27 ΔPDZ, or GFP alone as a control. Transfected cells were then fixed, and cell surface extope-CFTR was labeled using the external HA-tag to mark this pool for detection by laser-scanning cytometry (Fig. 5A). BHK extope-CFTR cells expressing SNX27 ΔPDZ presented 39% less CFTR on the cell surface compared with cells expressing WT SNX27 and 29% less than vector control. In contrast, BHK-extope-CFTR cells expressing WT SNX27 exhibited 17% more CFTR on the cell surface than did the control population (Fig. 5A).

**Fig. 5.**
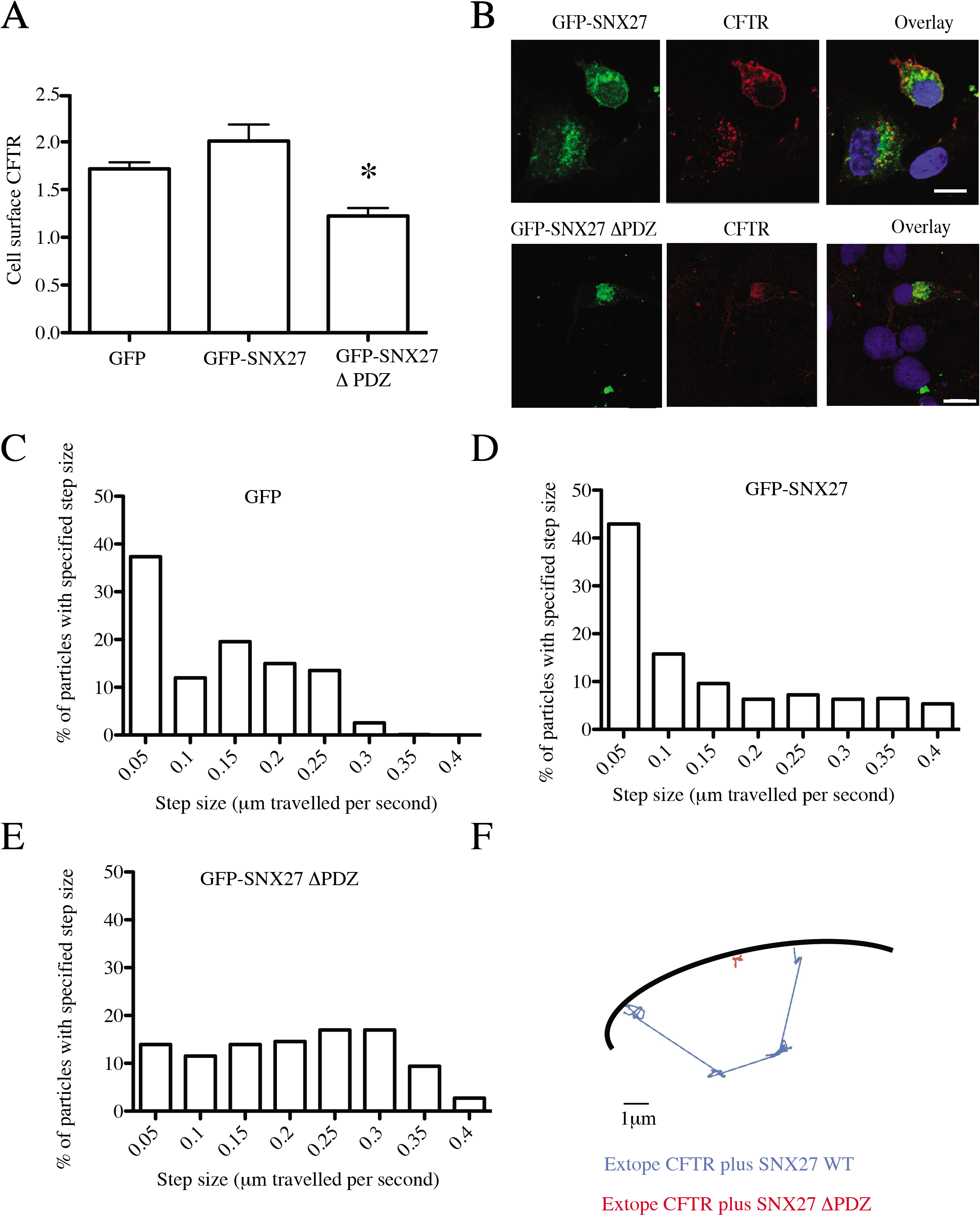
The SNX27 PDZ domain has an important role in CFTR trafficking. Stable BHK extope-CFTR cells were transiently transfected with GFP-tagged SNX27 wild type or SNX27 ΔPDZ. Cells were fixed and cell surface CFTR visualized using an antibody to the extracellular HA-tag. Laser scanning cytometry was used to quantify cell surface CFTR expression. At least 500 cells were counted per sample. Data was analyzed by students two-tailed t test. GFP-SNX27 ΔPDZ vs GFP control *P* < 0.05. GFP-SNX27 ΔPDZ vs GFP control *P* < 0.05. *n* = 3 are displayed as means +/- s.e.m. (A). Stable BHK extope-CFTR cells transiently transfected with GFP-tagged SNX27 wild type or GFP-SNX27 ΔPDZ were placed on ice and cell surface CFTR labeled with anti-HA antibody. The cells were subsequently restored to normal physiological conditions for 48hrs. Cells were fixed, permeabilized and incubated with secondary antibody to allow visualization of the CFTR protein. Internalized CFTR becomes trapped in a GFP-SNX27 ΔPDZ, but not wild type GFP-SNX27 positive compartment. Scale bar 10 μm (B). Extope CFTR cells were transfected with GFP; GFP-SNX27 or GFP-SNX27 ΔPDZ and labeled with anti-HA quantum dots on ice. Cells were restored to normal growth conditions and the quantum dot trajectories tracked at 1 frame per second (1HZ). Data is representative of at least 5 cells and 30 particles, in three separate experiments. Data is presented as the percentage of analyzed quantum dots that displayed a certain step size (lateral movement between frames) (C-E). Individual data for each of the three conditions are shown for GFP (C); GFP-SNX27 (D) and GFP-SNX27 DPDZ transfected cells (E). Typical trajectories illustrating the differences in CFTR-particle motility between cells expressing WT and mutant SNX27 (F).

SNX27 is demonstrated to play an essential role in PDZ-dependent trafficking of the beta(2)-adrenoreceptor (beta(2)AR) from endosomes to the plasma membrane. In that capacity, SNX27 functions as a cargo adaptor for loading beta (2)AR into the retromer complex, facilitating its sorting to the recycling pathway and away from the lysosomal degradation pathway (Lauffer et al., 2010; Temkin et al., 2011). To determine if cells expressing SNX27 ΔPDZ showed reduced recycling of cell-surface CFTR, BHK extope CFTR cells were transfected with WT GFP-SNX27 or GFP-SNX27 ΔPDZ expression constructs, and internalization of labeled CFTR from the cell surface was tracked by confocal microscopy over a 48 h time course. Despite similar rates of CFTR internalization in both populations of transfected cells, less cell surface CFTR was observed at 48 h in cells expressing GFP-SNX27 ΔPDZ. Furthermore, internalized CFTR became trapped in a SNX27 ΔPDZ positive compartment, suggesting impaired recycling back to the plasma membrane (Fig. 5B, bottom row). In contrast, most internalized CFTR in cells expressing WT GFP-SNX27 eventually moved to a SNX27-negative cellular compartment or returned to the plasma membrane (Fig. 5B, top row).

### Single particle tracking analyses of SNX27-mediated control of CFTR trafficking

Single particle tracking methods were used to further investigate the role of SNX27 in regulating CFTR recycling. In these experiments, BHK-extope-CFTR cells were transiently transfected with WT GFP-SNX27, GFP-SNX27 ΔPDZ, or GFP vector control expression cassettes, and cell-surface extope-CFTR was labeled with quantum dots which bound to the external extope HA-tag. Using a sampling window of 1 frame per second (1 Hz), movies were acquired to track labeled proteins over a period of 100 seconds. This relatively long sampling rate was chosen to distinguish interspersed periods of rapid long-range directional movement from more frequent short-range random Brownian motions. That is, CFTR-quantum dots undergoing Brownian motions moved very little, whereas quantum-dots being directionally transported through the endosomal system moved distinctly further from the origin (T0). Quantum dot trajectories were subsequently analyzed to compare lateral movements between individual frames. These movements report distances traveled by the particle in μm/second and define the “step size”.

Quantum dots bound to CFTR residing at the cell surface were expected to display small step sizes because of the confined diffusion imposed by PDZ-mediated surface anchorage (Haggie *et al*., 2006; Thelin *et al*., 2007). In contrast to cell surface CFTR, molecules undergoing active intracellular transport through the endosomal system would be recognized by their display of consistent directionality with increased step size. BHK extope-CFTR cells expressing GFP alone (Fig. 5C) or WT GFP-SNX27 (Fig. 5D) showed a high percentage of CFTR-quantum dots displaying small step sizes, consistent with most of the labeled CFTR remaining at the plasma membrane in these cells. CFTR-quantum dots which internalized from the cell surface rapidly recycled back to the plasma membrane and displayed increased step sizes as these molecules transited through the endosomal system. In contrast, BHK-CFTR cells expressing SNX27 ΔPDZ exhibited fewer quantum dots bound to the cell surface, consistent with our observations that overexpression of this mutant SNX27 reduced cell-surface CFTR levels. Moreover, internalized CFTR-particles became trapped within the endosomal system and transport back to the cell surface was not observed. These SNX27 ΔPDZ-expressing cells displayed an increase fraction of labeled particles moving with larger step sizes, and reciprocally decreased fractions of molecules showing low step sizes (Fig. 5E). Typical trajectories illustrating the differences in CFTR-particle motility between cells expressing WT and mutant SNX27 are shown in Fig. 5F.

### SNX27 regulates CFTR trafficking and protein levels in airway epithelia

The previous experiments all interrogated the behavior of ectopically expressed CFTR and SNX27 species. We therefore extended our investigation to the contribution of SNX27 in control of endogenous CFTR in polarized human airway epithelial Calu-3 cells. We had previously observed that CFTR localizes both to the apical surface of polarized Calu-3 cells and to sub-apical plasma membrane vesicles. Using the CFTR-interactor NHERF-1 as marker for the apical cell surface, we observed SNX27 in a sub-apical localization where it co-localized with the early endosome markers EEA1 and Rab5 (Fig. 6A). Staining of polarized Calu-3 cells with specific antibodies directed against SNX27 and CFTR revealed substantial co-localization of the two proteins directly below the apical surface (Figs 6B, C). These Calu-3 cell data report that the intracellular trafficking itineraries of CFTR and SNX27 intersect in endosomal compartments, and are congruent with the findings described above in non-airway cells.

**Fig. 6.**
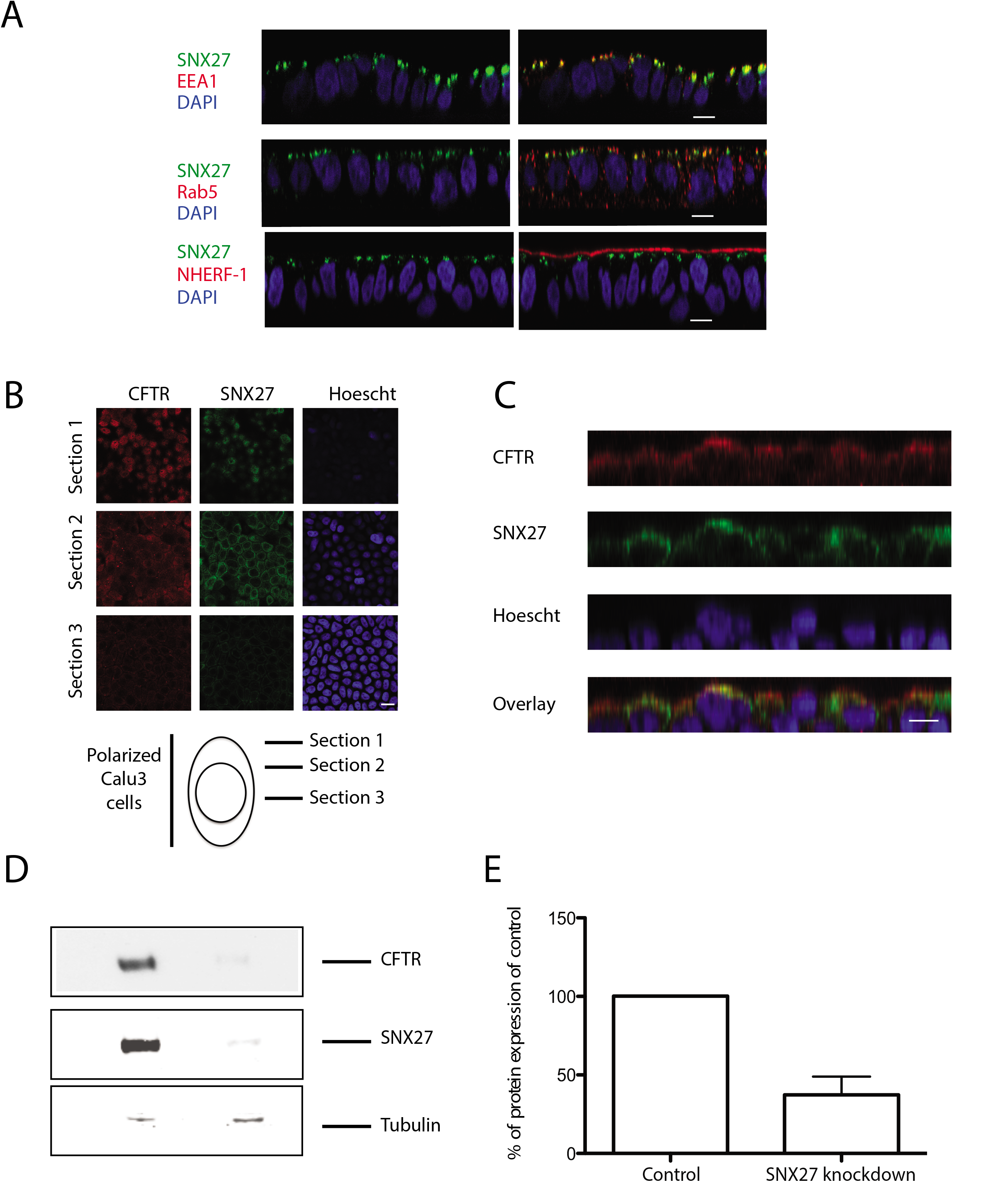
SNX27 localizes to subapical sites in polarized Calu-3 cells. Calu-3 cells were grown on permeable supports until transepithelial resistance reached 300 Ωcm^2^. Cells were fixed, permeabilized and stained with the indicated antibodies. Confocal slices revealed the expected localization of NHERF-1 at the apical membrane with SNX27 co-localizing with endosomal markers in a subapical sites (A). SNX27 co-localizes with intracellular pools of CFTR in polarized Calu-3 cells (B and C) Scale bars, 10 μm. Knockdown of CFTR in Calu-3 cells, results in a decrease in CFTR protein levels (D and E).

To address the role of SNX27 on CFTR trafficking in Calu-3 cells, lentiviruses driving inducible expression of either SNX27-directed or control shRNA were used to infect these airway epithelial cells. Successfully transduced cells were selected by a cell sorting regime that gated on the shRNA fluorescent tag, and efficacy of SNX27 knockdown was assessed 72 h post induction by immunoblotting. LI-COR^TM^ imaging showed 80% knockdown of SNX27 protein compared to control (Fig. 6D, middle panel). The knockdown cells showed strong (ca. 63%) reductions in endogenous CFTR levels (Fig. 6D, top panel; Fig. 6E), despite the fact that SNX27 knockdown in CFTR overexpressing cells were not observed to show similar reductions (data not shown). Thus, endogenous SNX27 is a major contributor to control of endogenous CFTR trafficking and homeostasis in human airway epithelia.

## Discussion

Protein-protein interactions between PDZ domains and PDZ-binding motifs localize receptors, ion channels, and many other types of proteins to specific regions of the cell. Localization of CFTR to the apical plasma membrane is essential for channel functionality and has been shown to require the PDZ-binding motif through studies using a mutant version of the protein lacking this region (CFTR-ΔTRL) (Swiatecka-Urban et al., 2002). CFTR-ΔTRL exhibited post-synthesis targeting to the apical membrane and its endocytosis was seemingly unaffected. However, the half-life of the channel was reduced, as was recycling of the internalized channel back to the cell surface (Swiatecka-Urban et al., 2002). The mechanism underlying the change in recycling efficiency of CFTR-ΔTRL has, as yet, not been elucidated.

Interactions between PDZ domain containing proteins and CFTR are known to give rise to compartment-specific regulation of the channel. The NHERF family of proteins localize to the sub-apical surface of polarized epithelial cells and interact with CFTR at the apical plasma membrane to: (1) anchor CFTR channels to the underlying cytoskeleton via ezrin/actin binding; (2) stabilize CFTR at the cell surface, and (3) compartmentalize CFTR with additional signaling proteins, such as regulatory kinases (Moyer et al., 1998; Short et al., 1998; Swiatecka-Urban et al., 2002). In contrast, overexpression of the PDZ domain containing protein GOPC, which localizes within the Golgi, reduces CFTR at the plasma membrane and increases its lysosomal-mediated degradation (Cheng et al., 2002; Cheng et al., 2004). The effect of GOPC on CFTR may be counteracted by NHERF-1, which competes for the CFTR C-terminus with GOPC to restore localization of CFTR at the cell surface (Cheng et al., 2002; Cheng et al., 2004). Our understanding of the regulation of CFTR trafficking by GOPC is further complicated by the recent identification of syntaxin 6 as a GOPC-interacting protein (Cheng *et al*., 2010). Silencing of syntaxin 6 increases wild type surface CFTR, but not CFTR lacking the C-terminal GOPC interaction site (Cheng et al., 2010). Recently a PDZ mediated interaction was discovered between CFTR and another Golgi protein, Golgi reassembly stacking protein 55 (Grasp55). In complete contrast to the interaction with GOPC transgenic expression of Grasp55 facilitated unconventional secretion to the plasma membrane of both wild type and ΔF508 CFTR, increasing their levels at the cell surface (Gee et al., 2011).

In this study, we identify SNX27 as novel binding partner of CFTR and show that binding is mediated by an interaction between the PDZ domain of SNX27 and the C-terminal PDZ-binding motif of CFTR. SNX27 belongs to a family of 33 PX domain-containing proteins but is unique in containing an N-terminal PDZ domain (Cullen 2008; Rincón et al., 2011). SNX27 is also the only sorting nexin family member with a FERM– like domain and has recently been shown to bind H-Ras through this region (Ghai et al., 2011). Since the identification of SNX1, discovered to bind to the C-terminus of the epidermal growth factor receptor (EGFR), sorting nexins have been strongly implicated in the endocytic trafficking of cell surface proteins (Cullen, 2008; Kurten et al., 1996; Worby and Dixon, 2002). SNX27 along with SNX17 and SNX31 belongs to a subfamily of early endosome localized SNX proteins, which promote retrieval and recycling of specific cargo populations (Cullen et al., 2008; Steinberg et al., 2013).

SNX27 was originally identified in rat neurons as the methamphetamine (MAP) inducible gene *Mrt1* (MAP responsive transcript 1) (Kajii et al., 2003). SNX27/Mrt1 was subsequently found to interact with the 5-hydroxytryptamine type 4 receptor variant a (5-HT4_(a)_R) and to target the receptor to the early endosome through a PDZ-dependent mechanism (Joubert et al., 2004). The PDZ domain of SNX27 has also been observed to interact with diacylglycerol kinase ζ, the cytohesin associated scaffolding protein CASP, the Kir3 potassium channel, the β1/2-adrenonergic receptors and N-Methyl-d-aspartate (NMDA) receptor 2C (NR2C) (Lunn et al., 2007; Rincón et al., 2007; Lauffer et al., 2010; Cai et al., 2011; Rincón et al., 2011; Temkin et al., 2011). More recently, our laboratory has identified PDZ mediated interaction between SNX27 and the PAK interactive exchange factor (βpix) a guanine nucleotide exchange factor for the Rho family of small GTPases (Valdes et al., 2011; Steinberg et al., 2013) and zonula occludens-2 (ZO-2) (Zimmerman et al., 2013). Recently quantitative proteomics has been employed to identify the SNX27 interactome identifying many more binding partners including the GLUT1 glucose transporter and a number of metal ion and amino acid transporters and has demonstrated their reduced surface expression in SNX27 knockdown cells (Steinberg et al., 2013).

We believe that SNX27 is an important regulator of CFTR endocytic trafficking because: 1) SNX27 is located in the early and recycling endosomes, which are known locations of CFTR trafficking; 2) SNX27 co-localized with internalized CFTR; 3) overexpression of SNX27 ΔPDZ or knockdown of endogenous SNX27 lead to decreased cell surface CFTR levels; 4) knockdown of endogenous SNX27 dramatically reduced overall CFTR protein levels.

Previous data has proposed diverse roles for SNX27 in the trafficking of interacting proteins, indicating that it may have distinct functions dependent on the protein it is trafficking (Joubert et al., 2004; Lauffer et al., 2010; Lunn et al., 2007). The interaction between SNX27 and the 5-hydroxytryptamine type 4 receptor (5-HT4R) or the Kir3 potassium channels proposed roles for SNX27 in promoting endocytosis or lysosomal delivery, respectively (Joubert et al., 2004; Lunn et al., 2007). Data for the beta(2)-adrenoreceptor (β2AR), in agreement with our observations for CFTR, demonstrate a role for SNX27 in PDZ mediated trafficking from the endosome to the plasma membrane (Lauffer et al., 2010; Temkin et al., 2011). We observed little colocalization of internalizing CFTR with SNX27 at early time points, and the rate of CFTR internalization was unaltered in cells over-expressing SNX27 or in SNX27 knockdown cells. In addition, co-localization and single particle trafficking data in cells over-expressing SNX27 ΔPDZ suggested that internalized CFTR becomes trapped in an endosomal compartment and does not recycle to the apical surface. Taken together, these data are consistent with a role in trafficking of CFTR through the endosomal recycling system.

How SNX27 regulates the endosomal trafficking of CFTR is unclear. One possibility is that SNX27 acts as an adaptor that facilitates CFTR binding to endosomes, stabilizing the internalized CFTR protein and chaperoning it through the endosomal system, thus preventing its degradation by the lysosome. Our observations are consistent with this hypothesis, since knockdown of SNX27 resulted in a dramatic reduction of over-expressed CFTR at the cell surface, and total endogenous CFTR levels. SNX27 has previously been reported as an essential adaptor of β-adrenergic receptors to retromer tubules where it interacted with the retromer-associated Wiskott-Aldrich syndrome protein and SCAR homologue (WASH) complex to promote PDZ-directed plasma membrane sorting. Knockdown of the retromer by RNAi inhibited β2AR recycling and redirected it to the lysosome (Temkin et al., 2011). Most recently SNX27 has been identified as a core component of a trafficking hub through independent association with both the WASH complex and the SNX-BAR-retromer (Steinberg et al., 2013). The mammalian SNX-BAR-retromer is composed of SNX1-SNX2 and SNX5-6 heterodimers loosely associating with a stable trimer of VPS26-29-35 and has a well-defined role in retrieval of endosome cargoes to the TGN (Arighi et al., 2004; Carlton et al., 2004; Seaman et al., 2004; Wassmer et al., 2007; 2009). The association of SNX27 with this complex suggests an additional role in mediating endosome-plasma membrane recycling (Temkin et al., 2011; Steinberg et al., 2013). Recent data consistent with this hypothesis demonstrate direct interaction between the SNX27 PDZ domain and VPS26 is both necessary and sufficient to prevent lysosomal entry of SNX27 cargo, while the SNX27 FERM-domain mediates interaction with SNX1 and possible SNX2 (Steinberg et al., 2013). It is highly possible that SNX27 acts in this manner to traffic CFTR, and our data fit this model. Intriguingly, CFTR has previously been shown to exist as part of a macromolecular complex with β2-adrenergic receptor, linked to it by NHERF (Naren *et al.*, 2003).

Recycling of cell surface proteins such as CFTR facilitates repeated cycles of functional action and provides a checkpoint to remove degraded, damaged, or misfolded proteins (Ghosh and Maxfield, 1995; Mellman, 1996). Given the rate of internalization (3-4% per min) and its restricted translation (Prince et al., 1999; Swiatecka-urban et al 2002; 2004) efficient recycling of CFTR is essential to maintain cell surface protein levels and to maintain its slow rate of turnover (t1/2 14-18h) (sharma et al., 2004; Swiatecka-urban et al., 2002; Picciano et al., 2003; reviewed Barrière et al., 2011). The mechanisms underlying this process are likely to be complex in order to facilitate the correct temporal and specific removal of a vast number of proteins from the cell surface to determine their fate. SNX27 is emerging as a regulator of the trafficking of a number of PDZ binding motif containing proteins and may well be involved in the trafficking of many more. How such specificity is achieved remains to be resolved, with specificity of action possibly being determined by the trafficked protein and distinct sets of regulators. One attractive possibility is that this occurs through regulation by GTPases, interacting with SNX27 through its C-terminal FERM-like domain.

The factors that regulate the surface expression of CFTR provide a mechanistic understanding of the underlying molecular defect in cystic fibrosis and are critical for the development and evaluation of therapeutic strategies aimed at correcting mutant CFTR proteins. The most common disease causing CFTR mutation, ΔF508, disrupts channel biosynthesis, resulting in a protein that is retained in the ER and does not traffic to the cell surface. A reduction in temperature or treatment with small-molecule correctors are two possibilities for enabling ΔF508 to escape the ER and reach the apical cell surface (Becq, 2010); however, the rescued mutant protein displays a defect in recycling (Cholon et al., 2010; Gentzsch et al., 2004; Sharma et al., 2004). While these methods are valuable for their ability to increase CFTR Cl^-^ currents, the durability of the therapeutic benefit will depend on the retention of CFTR on the cell surface. Thus, understanding the factors that regulate CFTR surface expression and recycling are of extreme importance in designing optimum treatment modalities.

## Acknowledgments

We would like to acknowledge Sharon L. Milgram (NIH, Bethesda MD 20892) for her input and guidance during early stages of the project, the help of Stephen M. Wincovitch of the National Human Genome Research Institute (NHGRI) microscopy core and Mark Ryherd and Stacie Anderson of the NHGRI flow cytometry core, NIH, Bethesda MD 20892. We would also like to thank Carol Parker and Ken Jacobson, UNC Chapel Hill, Chapel Hill, NC, 27599 and Li Shen Loo, Membrane and Biology Laboratory, Institute of Molecular and Cell Biology, Singapore.

---

**Supplemental Table 1.**
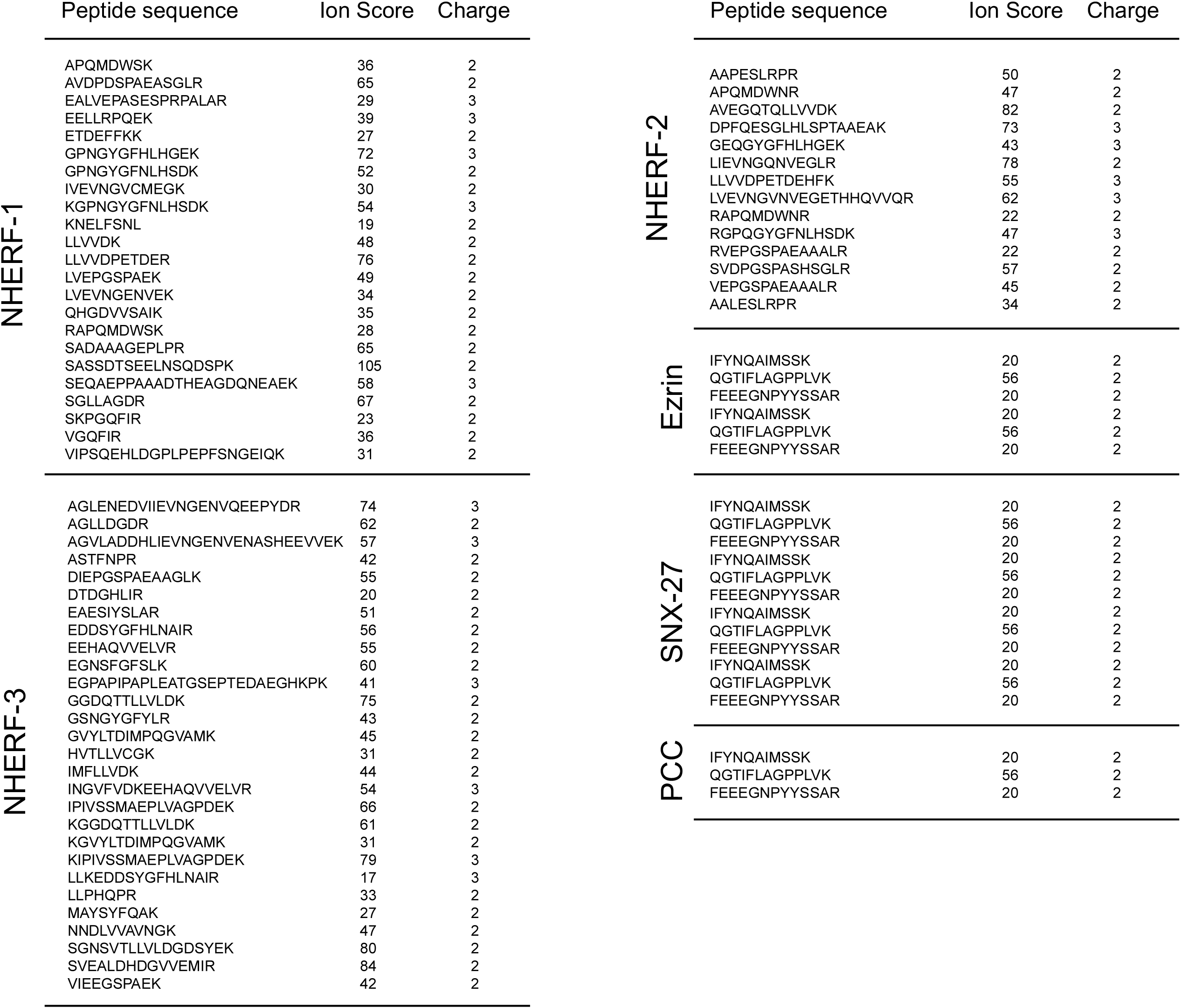

## References

Balch W. E. and Yates J. R., 3rd. (2011). Application of mass spectrometry to study proteomics and interactomics in cystic fibrosis. Methods Mol Biol 742, 227–47.

Barriere, H., Apaja, P., Okiyoneda, T. and Lukacs, G. L. (2011). Endocytic sorting of CFTR variants monitored by single-cell fluorescence ratiometric image analysis (FRIA) in living cells. Methods Mol Biol 741, 301–17.

Bates, I. R., Hebert, B., Luo, Y., Liao, J., Bachir, A. I., Kolin, D. L., Wiseman, P. W. and Hanrahan, J. W. (2006). Membrane lateral diffusion and capture of CFTR within transient confinement zones. Biophys J 91, 1046–58.

Becq F. (2010). Cystic fibrosis transmembrane conductance regulator modulators for personalized drug treatment of cystic fibrosis: progress to date. Drugs 70, 241–59.

Benharouga, M., Sharma, M., So, J., Haardt, M., Drzymala, L., Popov, M., Schwapach, B., Grinstein, S., Du, K. and Lukacs, G. L. (2003). The role of the C terminus and Na+/H+ exchanger regulatory factor in the functional expression of cystic fibrosis transmembrane conductance regulator in nonpolarized cells and epithelia. J Biol Chem 278, 22079–89.

Brown, C. R., Hong-Brown, L. Q., Biwersi, J., Verkman, A. S. and Welch, W. J. (1996). Chemical chaperones correct the mutant phenotype of the delta F508 cystic fibrosis transmembrane conductance regulator protein. Cell Stress Chaperones 1, 117–25.

Cai, L., Loo, L. S., Atlashkin, V., Hanson, B. J. and Hong, W. (2011). Deficiency of sorting nexin 27 (SNX27) leads to growth retardation and elevated levels of N-methyl-D-aspartate receptor 2C (NR2C). Mol Cell Biol 31, 1734–47.

Chang, X. B., Cui, L., Hou, Y. X., Jensen, T. J., Aleksandrov, A. A., Mengos, A. and Riordan, J. R. (1999). Removal of multiple arginine-framed trafficking signals overcomes misprocessing of delta F508 CFTR present in most patients with cystic fibrosis. Mol Cell 4, 137–42.

Chen, Y., Veracini, L., Benistant, C. and Jacobson, K. (2009). The transmembrane protein CBP plays a role in transiently anchoring small clusters of Thy-1, a GPI-anchored protein, to the cytoskeleton. J Cell Sci 122, 3966–72.

Cheng, J., Cebotaru, V., Cebotaru, L. and Guggino, W. B. (2010). Syntaxin 6 and CAL mediate the degradation of the cystic fibrosis transmembrane conductance regulator. Mol Biol Cell 21, 1178–87.

Cheng, J., Moyer, B. D., Milewski, M., Loffing, J., Ikeda, M., Mickle, J. E., Cutting, G. R., Li, M., Stanton, B. A. and Guggino, W. B. (2002). A Golgi-associated PDZ domain protein modulates cystic fibrosis transmembrane regulator plasma membrane expression. J Biol Chem 277, 3520–9.

Cheng, J., Wang, H. and Guggino, W. B. (2004). Modulation of mature cystic fibrosis transmembrane regulator protein by the PDZ domain protein CAL. J Biol Chem 279, 1892–8.

Cholon, D. M., O’Neal, W. K., Randell, S. H., Riordan, J. R. and Gentzsch, M. (2010). Modulation of endocytic trafficking and apical stability of CFTR in primary human airway epithelial cultures. Am J Physiol Lung Cell Mol Physiol 298, L304–14.

Cullen P. J. (2008). Endosomal sorting and signalling: an emerging role for sorting nexins. Nat Rev Mol Cell Biol 9, 574–82.

Denning, G. M., Anderson, M. P., Amara, J. F., Marshall, J., Smith, A. E. and Welsh, M. J. (1992). Processing of mutant cystic fibrosis transmembrane conductance regulator is temperature-sensitive. Nature 358, 761–4.

Egan, M. E., Pearson, M., Weiner, S. A., Rajendran, V., Rubin, D., Glockner-Pagel, J., Canny, S., Du, K., Lukacs, G. L. and Caplan, M. J. (2004). Curcumin, a major constituent of turmeric, corrects cystic fibrosis defects. Science 304, 600–2.

Gee, H. Y., Noh, S. H., Tang, B. L., Kim, K. H. and Lee, M. G. (2011). Rescue of DeltaF508-CFTR Trafficking via a GRASP-Dependent Unconventional Secretion Pathway. Cell 146, 746–60.

Gee, H. Y., Tang, B. L., Kim, K. H. and Lee, M. G. (2010). Syntaxin 16 binds to cystic fibrosis transmembrane conductance regulator and regulates its membrane trafficking in epithelial cells. J Biol Chem 285, 35519–27.

Gentzsch, M., Chang, X. B., Cui, L., Wu, Y., Ozols, V. V., Choudhury, A., Pagano, R. E. and Riordan, J. R. (2004). Endocytic trafficking routes of wild type and DeltaF508 cystic fibrosis transmembrane conductance regulator. Mol Biol Cell 15, 268496.

Ghosh R. N. and Maxfield F. R. (1995). Evidence for nonvectorial, retrograde transferrin trafficking in the early endosomes of HEp2 cells. J Cell Biol 128, 549–61.

Grubb, B. R., Gabriel, S. E., Mengos, A., Gentzsch, M., Randell, S. H., Van Heeckeren, A. M., Knowles, M. R., Drumm, M. L., Riordan, J. R. and Boucher, R. C. (2006). SERCA pump inhibitors do not correct biosynthetic arrest of deltaF508 CFTR in cystic fibrosis. Am JRespir Cell Mol Biol 34, 355–63.

Haggie, P. M., Kim, J. K., Lukacs, G. L. and Verkman, A. S. (2006). Tracking of quantum dot-labeled CFTR shows near immobilization by C-terminal PDZ interactions. Mol Biol Cell 17, 4937–45.

Haggie, P. M., Stanton, B. A. and Verkman, A. S. (2004). Increased diffusional mobility of CFTR at the plasma membrane after deletion of its C-terminal PDZ binding motif. J Biol Chem 279, 5494–500.

Han, W., Kim, K. H., Jo, M. J., Lee, J. H., Yang, J., Doctor, R. B., Moe, O. W., Lee, J., Kim, E. and Lee, M. G. (2006). Shank2 associates with and regulates Na+/H+ exchanger 3. J Biol Chem 281, 1461–9.

Hayashi, H., Naoi, S., Nakagawa, T., Nishikawa, T., Fukuda, H., Imajoh-Ohmi, S., Kondo, A., Kubo, K., Yabuki, T., Hattori, A. et al. (2012). Sorting Nexin 27 Interacts with Multidrug Resistance-associated Protein 4 (MRP4) and Mediates Internalization of MRP4. J Biol Chem 287, 15054–65.

Joubert, L., Hanson, B., Barthet, G., Sebben, M., Claeysen, S., Hong, W., Marin, P., Dumuis, A. and Bockaert, J. (2004). New sorting nexin (SNX27) and NHERF specifically interact with the 5-HT4a receptor splice variant: roles in receptor targeting. J Cell Sci 117, 5367–79.

Kajii, Y., Muraoka, S., Hiraoka, S., Fujiyama, K., Umino, A. and Nishikawa, T. (2003). A developmentally regulated and psychostimulant-inducible novel rat gene mrt1 encoding PDZ-PX proteins isolated in the neocortex. Mol Psychiatry 8, 434–44.

Kim, J. Y., Han, W., Namkung, W., Lee, J. H., Kim, K. H., Shin, H., Kim, E. and Lee, M. G. (2004). Inhibitory regulation of cystic fibrosis transmembrane conductance regulator anion-transporting activities by Shank2. J Biol Chem 279, 10389–96.

Koulov A. V., Lapointe P., Lu B., Razvi A., Coppinger J., Dong M. Q., Matteson J., Laister R., Arrowsmith C., Yates J. R., 3rd et al. (2010). Biological and structural basis for Aha1 regulation of Hsp90 ATPase activity in maintaining proteostasis in the human disease cystic fibrosis. Mol Biol Cell 21, 871–84.

Kulkarni, R. P., Castelino, K., Majumdar, A. and Fraser, S. E. (2006). Intracellular transport dynamics of endosomes containing DNA polyplexes along the microtubule network. Biophysical Journal 90, L42–L44.

Kurten, R. C., Cadena, D. L. and Gill, G. N. (1996). Enhanced degradation of EGF receptors by a sorting nexin, SNX1. Science 272, 1008–10.

Lauffer, B. E., Melero, C., Temkin, P., Lei, C., Hong, W., Kortemme, T. and von Zastrow, M. (2010). SNX27 mediates PDZ-directed sorting from endosomes to the plasma membrane. J Cell Biol 190, 565–74.

Lee, J. H., Richter, W., Namkung, W., Kim, K. H., Kim, E., Conti, M. and Lee, M. G. (2007). Dynamic regulation of cystic fibrosis transmembrane conductance regulator by competitive interactions of molecular adaptors. J Biol Chem 282, 10414–22.

Lee, J. S., Kim, J. H., Jang, I. H., Kim, H. S., Han, J. M., Kazlauskas, A., Yagisawa, H., Suh, P. G. and Ryu, S. H. (2005). Phosphatidylinositol (3,4,5)-trisphosphate specifically interacts with the phox homology domain of phospholipase D1 and stimulates its activity. Journal of Cell Science 118, 4405–4413.

Lukacs, G. L., Mohamed, A., Kartner, N., Chang, X. B., Riordan, J. R. and Grinstein, S. (1994). Conformational maturation of CFTR but not its mutant counterpart (delta F508) occurs in the endoplasmic reticulum and requires ATP. EMBO J 13, 6076–86.

Lunn M. L., Nassirpour R., Arrabit C., Tan J., McLeod I., Arias C. M., Sawchenko P. E., Yates J. R., 3rd and Slesinger, P. A. (2007). A unique sorting nexin regulates trafficking of potassium channels via a PDZ domain interaction. Nat Neurosci 10, 1249–59.

Mellman I. (1996). Endocytosis and molecular sorting. Annu Rev Cell Dev Biol 12, 575–625.

Moyer, B. D., Denton, J., Karlson, K. H., Reynolds, D., Wang, S., Mickle, J. E., Milewski, M., Cutting, G. R., Guggino, W. B., Li, M. et al. (1999). A PDZ-interacting domain in CFTR is an apical membrane polarization signal. J Clin Invest 104, 1353–61.

Moyer, B. D., Loffing, J., Schwiebert, E. M., Loffing-Cueni, D., Halpin, P. A., Karlson, K. H., IsmailovII, Guggino, W. B., Langford, G. M. and Stanton, B. A. (1998). Membrane trafficking of the cystic fibrosis gene product, cystic fibrosis transmembrane conductance regulator, tagged with green fluorescent protein in madin-darby canine kidney cells. J Biol Chem 273, 21759–68.

Murray, J. W., Bananis, E. and Wolkoff, A. W. (2000). Reconstitution of ATP-dependent movement of endocytic vesicles along microtubules in vitro: an oscillatory bidirectional process. Mol Biol Cell 11, 419–33.

Picciano, J. A., Ameen, N., Grant, B. D. and Bradbury, N. A. (2003). Rme-1 regulates the recycling of the cystic fibrosis transmembrane conductance regulator. Am J Physiol Cell Physiol 285, C1009–18.

Raghuram, V., Mak, D. O. and Foskett, J. K. (2001). Regulation of cystic fibrosis transmembrane conductance regulator single-channel gating by bivalent PDZ-domain-mediated interaction. Proc Natl Acad Sci U S A 98, 1300–5.

Rincon, E., Saez de Guinoa, J., Gharbi, S. I., Sorzano, C. O., Carrasco, Y. R. and Merida, I. (2011). Translocation dynamics of sorting nexin 27 in activated T cells. J Cell Sci 124, 776–88.

Rincon, E., Santos, T., Avila-Flores, A., Albar, J. P., Lalioti, V., Lei, C., Hong, W. and Merida, I. (2007). Proteomics identification of sorting nexin 27 as a diacylglycerol kinase zeta-associated protein: new diacylglycerol kinase roles in endocytic recycling. Mol Cell Proteomics 6, 1073–87.

Riordan J. R. (2008). CFTR function and prospects for therapy. Annu Rev Biochem 77, 701–26.

Riordan, J. R., Rommens, J. M., Kerem, B., Alon, N., Rozmahel, R., Grzelczak, Z., Zielenski, J., Lok, S., Plavsic, N., Chou, J. L. et al. (1989). Identification of the cystic fibrosis gene: cloning and characterization of complementary DNA. Science 245, 1066–73.

Sharma, M., Pampinella, F., Nemes, C., Benharouga, M., So, J., Du, K., Bache, K. G., Papsin, B., Zerangue, N., Stenmark, H. et al. (2004). Misfolding diverts CFTR from recycling to degradation: quality control at early endosomes. J Cell Biol 164, 923–33.

Short, D. B., Trotter, K. W., Reczek, D., Kreda, S. M., Bretscher, A., Boucher, R. C., Stutts, M. J. and Milgram, S. L. (1998). An apical PDZ protein anchors the cystic fibrosis transmembrane conductance regulator to the cytoskeleton. J Biol Chem 273, 19797–801.

Silvis, M. R., Bertrand, C. A., Ameen, N., Golin-Bisello, F., Butterworth M. B., Frizzell, R. A. and Bradbury, N. A. (2009). Rab11b regulates the apical recycling of the cystic fibrosis transmembrane conductance regulator in polarized intestinal epithelial cells. Mol Biol Cell 20, 2337–50.

Stahelin, R. V., Burian, A., Bruzik, K. S., Murray, D. and Cho, W. W. (2003). Membrane binding mechanisms of the PX domains of NADPH oxidase p40(phox) and p47(phox). Journal of Biological Chemistry 278, 14469–14479.

Sun, F., Hug, M. J., Lewarchik, C. M., Yun, C. H., Bradbury, N. A. and Frizzell, R. A. (2000). E3KARP mediates the association of ezrin and protein kinase A with the cystic fibrosis transmembrane conductance regulator in airway cells. J Biol Chem 275, 29539–46.

Swiatecka-Urban, A., Boyd, C., Coutermarsh, B., Karlson, K. H., Barnaby, R., Aschenbrenner, L., Langford, G. M., Hasson, T. and Stanton, B. A. (2004). Myosin VI regulates endocytosis of the cystic fibrosis transmembrane conductance regulator. J Biol Chem 279, 38025–31.

Swiatecka-Urban, A., Brown, A., Moreau-Marquis, S., Renuka, J., Coutermarsh, B., Barnaby, R., Karlson, K. H., Flotte, T. R., Fukuda, M., Langford, G. M. et al. (2005). The short apical membrane half-life of rescued {Delta}F508-cystic fibrosis transmembrane conductance regulator (CFTR) results from accelerated endocytosis of {Delta}F508-CFTR in polarized human airway epithelial cells. J Biol Chem 280, 36762–72.

Swiatecka-Urban, A., Duhaime, M., Coutermarsh, B., Karlson, K. H., Collawn, J., Milewski, M., Cutting, G. R., Guggino, W. B., Langford, G. and Stanton, B. A. (2002). PDZ domain interaction controls the endocytic recycling of the cystic fibrosis transmembrane conductance regulator. J Biol Chem 277, 40099–105.

Temkin, P., Lauffer, B., Jager, S., Cimermancic, P., Krogan, N. J. and von Zastrow, M. (2011). SNX27 mediates retromer tubule entry and endosome-to-plasma membrane trafficking of signalling receptors. Nature Cell Biology 13, 715–U199.

Thelin, W. R., Chen, Y., Gentzsch, M., Kreda, S. M., Sallee, J. L., Scarlett, C. O., Borchers, C. H., Jacobson, K., Stutts, M. J. and Milgram, S. L. (2007). Direct interaction with filamins modulates the stability and plasma membrane expression of CFTR. J Clin Invest 117, 364–74.

Thelin, W. R., Kesimer, M., Tarran, R., Kreda, S. M., Grubb, B. R., Sheehan, J. K., Stutts, M. J. and Milgram, S. L. (2005). The cystic fibrosis transmembrane conductance regulator is regulated by a direct interaction with the protein phosphatase 2A. J Biol Chem 280, 41512–20.

Tsui L. C. (1992). The spectrum of cystic fibrosis mutations. Trends Genet 8, 392–8.

Valdes, J. L., Tang, J., McDermott, M. I., Kuo, J. C., Zimmerman, S. P., Wincovitch, S. M., Waterman, C. M., Milgram, S. L. and Playford, M. P. (2011). Sorting Nexin 27 regulates trafficking of a PAK interacting exchange factor ({beta}PIX)-G-protein-coupled receptor kinase interacting protein (GIT) complex via a PDZ domain interaction. J Biol Chem.

van der Schaar, H. M., Rust, M. J., Chen, C., van der Ende-Metselaar, H., Wilschut, J., Zhuang, X. W. and Smit, J. M. (2008). Dissecting the Cell Entry Pathway of Dengue Virus by Single-Particle Tracking in Living Cells. Plos Pathogens 4.

van Weering, J. R. T., Verkade, P. and Cullen, P. J. (2012). SNX-BAR-Mediated Endosome Tubulation is Co-ordinated with Endosome Maturation. Traffic 13, 94–107.

Wang, S., Raab, R. W., Schatz, P. J., Guggino, W. B. and Li, M. (1998). Peptide binding consensus of the NHE-RF-PDZ1 domain matches the C-terminal sequence of cystic fibrosis transmembrane conductance regulator (CFTR). FEBS Lett 427, 103–8.

Wang, S., Yue, H., Derin, R. B., Guggino, W. B. and Li, M. (2000). Accessory protein facilitated CFTR-CFTR interaction, a molecular mechanism to potentiate the chloride channel activity. Cell 103, 169–79.

Wang, X., Venable, J., LaPointe, P., Hutt, D. M., Koulov, A. V., Coppinger, J., Gurkan, C., Kellner, W., Matteson, J., Plutner, H. et al. (2006). Hsp90 cochaperone Aha1 downregulation rescues misfolding of CFTR in cystic fibrosis. Cell 127, 803–15.

Ward C. L. and Kopito R. R. (1994). Intracellular turnover of cystic fibrosis transmembrane conductance regulator. Inefficient processing and rapid degradation of wild-type and mutant proteins. J Biol Chem 269, 25710–8.

Weinman, E. J., Hall, R. A., Friedman, P. A., Liu-Chen, L. Y. and Shenolikar, S. (2006). The association of NHERF adaptor proteins with g protein-coupled receptors and receptor tyrosine kinases. Annu Rev Physiol 68, 491–505.

Wood H. G. and Barden R. E. (1977). Biotin Enzymes. Annual Review of Biochemistry 46, 385–413.

Worby C. A. and Dixon J. E. (2002). Sorting out the cellular functions of sorting nexins. Nat Rev Mol Cell Biol 3, 919–31.

